# Host specificity of gastrointestinal parasites in free-ranging sloths from Costa Rica

**DOI:** 10.1101/2024.11.22.624753

**Authors:** Ezequiel A. Vanderhoeven, Madeleine G. Florida, Rebecca Cliffe, José Guzman, Juliana Notarnicola, Tyler R. Kartzinel

## Abstract

The diversity and host specificity of gastrointestinal parasites infecting free-ranging sloths is poorly known. We characterized gastrointestinal parasites of two sloth species from Costa Rica— three-fingered sloth (*Bradypus variegatus*) and two-fingered sloth (*Choloepus hoffmanni*)—for the first time in both a primary forest and an urban habitat. We asked whether host-parasite interactions were predominantly structured by host identity, the habitats in which hosts occurred, or both. Protozoa and nematode eggs were present in both species, but cestode eggs were recorded only in *C. hoffmanni*. We found eight parasitic morphotypes, which matches the total number of parasite taxa described in sloths over the past 100 years, and these first reports of gastrointestinal parasites of sloths from primary forest radically expand our knowledge of the general diversity and host specificity of sloth parasites. We found no significant difference in overall parasite richness between sloth species or habitats, but the parasite richness of *C. hoffmanni* was 2-fold greater in the primary forest vs. urban habitat. As no parasite sharing was observed between sloth species, we found strong and significant differences in parasite composition between host species regardless of habitat. In *B. variegatus*, we observed eggs of four nematode taxa (Spirocercidae, Subuluroidea, Spirurida, Ascaridida) and cysts of Eimeriidae (Apicomplexa). By contrast, in *C. hoffmanni*, we observed cestodes (Anoplocephalidae), a different nematode from the family Spirocercidae and also cysts of Eimeriidae (Apicomplexa). Many rare taxa were recorded only in samples from the primary forest, and these did not match any sloth parasites that had been previously described in the literature, suggesting that at least some could be undescribed species. Together, these results highlight the need for further research in comparative wildlife parasitology, the characterization of host-parasite transmission networks, and the identification of any intermediate hosts that may be relevant to sloth health.

## Introduction

Insights from disease ecology research have repeatedly shown that host susceptibility to parasites can be strongly influenced by the ecological context in which each host population occurs, and that the communities of parasites to which these hosts are exposed can have a profound influence on their health and survival (Preston and Johnson 2010; Irvine, 2006; Wood, 2023). Theoretically, the parasite assemblage associated with co-occurring host species from divergent mammalian lineages could be structured primarily by host phylogeny and any associated functional differences in anatomy, immunity, and/or behavioral ecology (Morand, 2015). Yet, although the study of gastrointestinal (GI) parasites is common in domestic animals, for which taxonomic keys facilitate identification, there is a scarcity of even coprological research into the GI parasites of wildlife mammals (Kaufmann, 2013; Thienpont et al. 1979; Agostini et al. 2018; Solórzano-García & de Leon 2017; de Almeida Curi et al. 2010).

Two-and three-toed sloths are distantly related members of the Pilosa (Xenarthra) order. Each sloth lineage evolved anatomical and behavioral characteristics associated with their slow, herbivorous, arboreal lifestyles from a terrestrial common ancestor via convergent evolution (Chiarello, 1998; Gaudin, 2003; Toledo et al. 2015 Hayssen, 2010, 2011; Urbani & Bosque, 2007). Fossil sloths were diverse and geographically widespread, extending in the southernmost Chile, Argentine Patagonia, and possibly Antarctica in the south and to the south of Alaska in the north (Toledo et al. 2015). Two-and three-toed sloth lineages diverged ∼30 million years ago, and since the megafaunal extinction of the terminal Pleistocene they have maintained broadly overlapping ranges across Central and South America (Toledo et al. 2015). Sloths are keystone herbivores of the Neotropical forests. Today, populations occur in a diversity of habitats including forests, agroecosystems, and urban environments where they may often come into close contact with humans or domestic animals—creating opportunities for GI parasite transmission (Brandão et al. 2019; Hayssen, 2010, 2011; Smith & Ruple, 2021; Superina et al. 2010; Superina & Loughry, 2015). Yet most prior studies of sloth GI parasites have involved captive animals, with only a few reports of necropsies involving wild animals (Dinis & Oliveira, 1999; Sibaja-Morales et al., 2009; Michel et al. 2017; Araujo et al. 2021). Studies of wild sloth parasites have tended instead to focus on their potential role as reservoirs for *Leishmania braziliensis* and other blood parasites such as *Trypanosoma* spp. and microfilariae that pose risks to humans (Gilmore et al. 2001; Herrer & Christensen, 1980; Lainson et al. 1989; Muñoz-García et al. 2019; Sant’Ana et al. 2020). Knowledge of GI parasites circulating in wild sloth populations is thus a priority at the intersection of wildlife ecology, conservation, and public health (Carlson et al. 2020; Pedersen & Fenton, 2007).

In Costa Rica, two species of sloths are distributed across a range of environmental conditions: Hoffmann’s two-toed sloths (*Choloepus hoffmanni* Peters,1858) and the brown-throated three-toed sloths (*Bradypus variegatus* Schinz, 1825; Santos et al. 2019; Superina et al., 2010). Given their broadly overlapping ecologies and their many convergent anatomical and behavioral characteristics—especially in their unique digestive morphophysiologies—wild populations of these co-occurring species in different habitats provide unique opportunities to evaluate the extent to which GI parasite assemblages are structured primarily by host identity or their shared ecological conditions. Therefore, we tested the hypotheses that the diversity and composition of GI parasites communities infecting these two species of wild sloths are determined primarily by (1) host species identity, (2) local habitat type, or (3) that they are modulated by both. Results will address a critical research need concerning our basic understanding of host-parasite interactions in tropical wildlife, the ability to effectively monitor wildlife diseases and potential zoonoses, and the success of numerous conservation activities in the region.

## Materials and Methods

### Study sites

We collected samples during March-July 2023 across the Caribbean coast of Costa Rica (Figure 1). Principal study sites included La Selva Biological Station and the city of Puerto Viejo de Talamanca. La Selva is a field station operated by the Organization for Tropical Studies (OTS) in Sarapiquí of northeastern Costa Rica, comprising 1,536 hectares of protected lowland tropical rainforest (10° 25.32’ N, 84° 00.9’ W; elevation 37–150 mASL;) adjoining the 47,000-hectare Braulio Carrillo National Park (Figure 1; Matlock & Hartshorn, 1999). Puerto Viejo de Talamanca (9.65°N, 82.75°W, elevation 0–4 mASL) is a small, but densely urbanized coastal town in southeastern Costa Rica, near both Cahuita and Manzanillo National Parks (Figure 1; Lindshield, 2016; Emard & Nelson, 2021; Fan & Lindshield, 2022).

**Figure 1.**
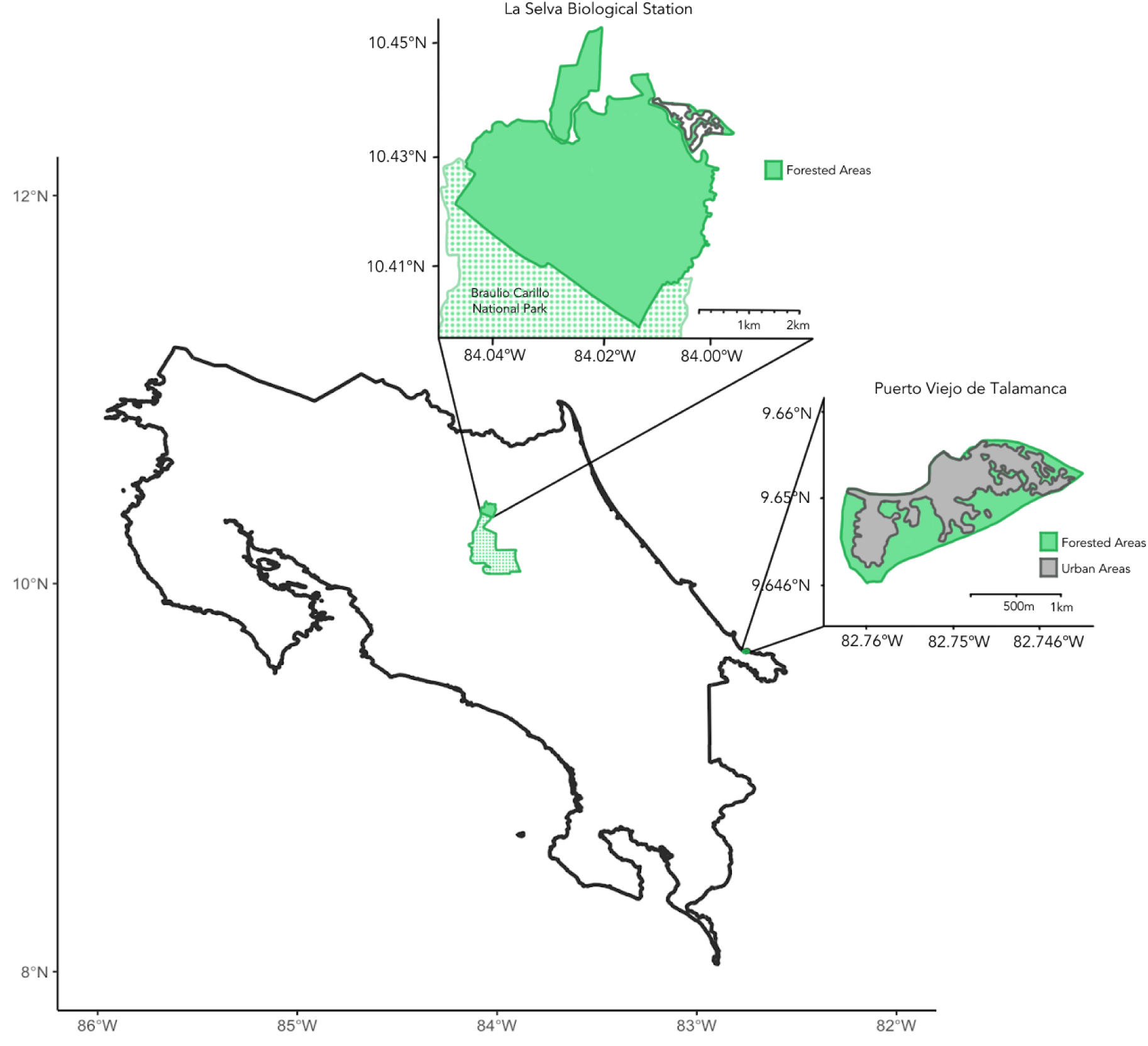
Map of Costa Rica, with insets showing the two main sites of sample collection, including La Selva Biological Station to the northwest and the city of Puerto Viejo de Talamanca to the southeast. The green shading shows areas of tree cover and the dotted areas in the northern inset show Braulio Carillo National Park, which is contiguous with La Selva Biological Station.

### Sample collection

We obtained samples from sloth feces using two sampling methods. First, we conducted active searches on foot through trails and roads for signs of sloth feces. Fecal samples were found by searching the bases of trees used as latrines. Second, we received samples from veterinarians shortly after sloths were newly admitted to wildlife hospitals. Samples were stored in 5% formalin for parasitological analyses. Permits were provided by the National System of Conservation Areas of Costa Rica (SINAC-ACC-PI-LC-052-2 023).

### Parasitology techniques

To detect a wide variety of parasitic structures including helminth eggs, larvae, and protozoan cysts, we used a set of techniques involving concentration, sedimentation, and flotation. First, we used the modified Telemann method, which is a sedimentation and concentration technique that allows visualization of parasitic structures such as heavy and light eggs, cysts, and larvae, especially in samples that have high concentrations of neutral fats and free fatty acids (Thienpont et al. 1979). The procedure consisted of homogenizing 1-2 g of fecal material preserved in formalin with water (5-6 pellets from *B. variegatus*; 1-3 pellets from *C. hoffmanni*). This was filtered through a strainer over a funnel, and 10 ml of filtrate was collected in a 15-ml conical tube to which 2 ml of sulfuric/ethyl ether was added and shaken (stopping to vent halfway through and after shaking). The resulting mixture was centrifuged at 1,500 rpm for 5 min to pellet the parasitic elements before removing the supernatant. The pellet was placed on a slide for observation using a Pasteur pipette and stained with a drop of 1% Lugol’s solution for visualization. Second, a modified Sheather method involved the use of a supersaturated sugar solution view light helminth eggs and oocysts of *Cryptosporidium* spp. and *Giardia* spp. (Sheather, 1923). It began by homogenizing 1-2 g of fecal material in water, followed by filtration through a strainer placed over a funnel from which 10 ml of filtrate was collected in a 15 ml conical tube. The sample was centrifuged at 2,500 rpm for 2 min, the supernatant discarded, and the tube filled with Benbrok solution (density 1.300) to 1 cm from the rim. The tube was shaken until the sediment dissolved and was then centrifuged at 1,000 rpm for 2 min. More Benbrok solution was added until a dome held in place by surface tension formed above the lip of the tube. A coverslip placed on top of the tube was left to incubate for 10 min before observation on a microscope at 10x and 40x magnification. We used an AmScope B400 Series microscope and took pictures with an AmScope MU1400 CMOS C-Mount Microscope Camera with Reduction Lens. Parasite structures were measured using ImageJ and eggs were identified to the finest taxonomic resolution possible according to relevant taxonomic keys and specific literature on helminths from sloths (Kaufmann, 2013; Vicente et al. 1997; Anderson et al. 2000; Khalil et al. 1994).

### Statistical Analysis

First, we tested for statistically significant differences in mean parasite richness according to sloth species, habitat, and the sloth species × habitat interaction using ANOVA. We based this richness analysis on all 38 samples, including 13 that tested negative for parasites and therefore had richness scores of zero. Then, after excluding the 13 samples in which no parasites were identified, we quantified pairwise similarity among the 25 positive sloth samples using the Jaccard Dissimilarity Index (0 = completely overlapping parasites; 1 = completely different). We tested for significant differences in parasite composition according to sloth species, habitat, and the sloth species × habitat interaction using permutational MANOVA (PERMANOVA) with 999 permutations with the *adonis2* function in *vegan* (Jaccard, 1912; Oksanen et al. 2024). All statistical analyses were performed in R (R Core Development Team, 2023).

## Results

### Parasite identification and prevalence

We collected 22 fecal samples from *B. variegatus* and 16 from *C. hoffmanni*. Of the 22 *B. variegatus* samples, 12 were from primary forest and 10 were from urban habitat with 64% testing positive for parasites (50% disturbed, 50% undisturbed). Of the 16 *C. hoffmanni* samples, 5 were from undisturbed sites and 11 were from disturbed sites, with 69% testing positive (64% disturbed, 80% undisturbed). We classified parasite structures into eight morphotypes based on a thorough review of the relevant literature (Tables 1, S1; Appendix S1). In *B. variegatus*, we identified four distinct types of Nematoda eggs and protozoa of the family Eimeriidae (Apicomplexa; Table 1; Figure 2). In *C. hoffmanni*, we observed one morphotype of Nematoda, one morphotype of Anoplocephalidae (Cestoda), and cysts of Eimeriidae (Apicomplexa; Figure 3). In both host species, Eimeriidae cysts showed the highest prevalence (Table 1).

**Figure 2.**
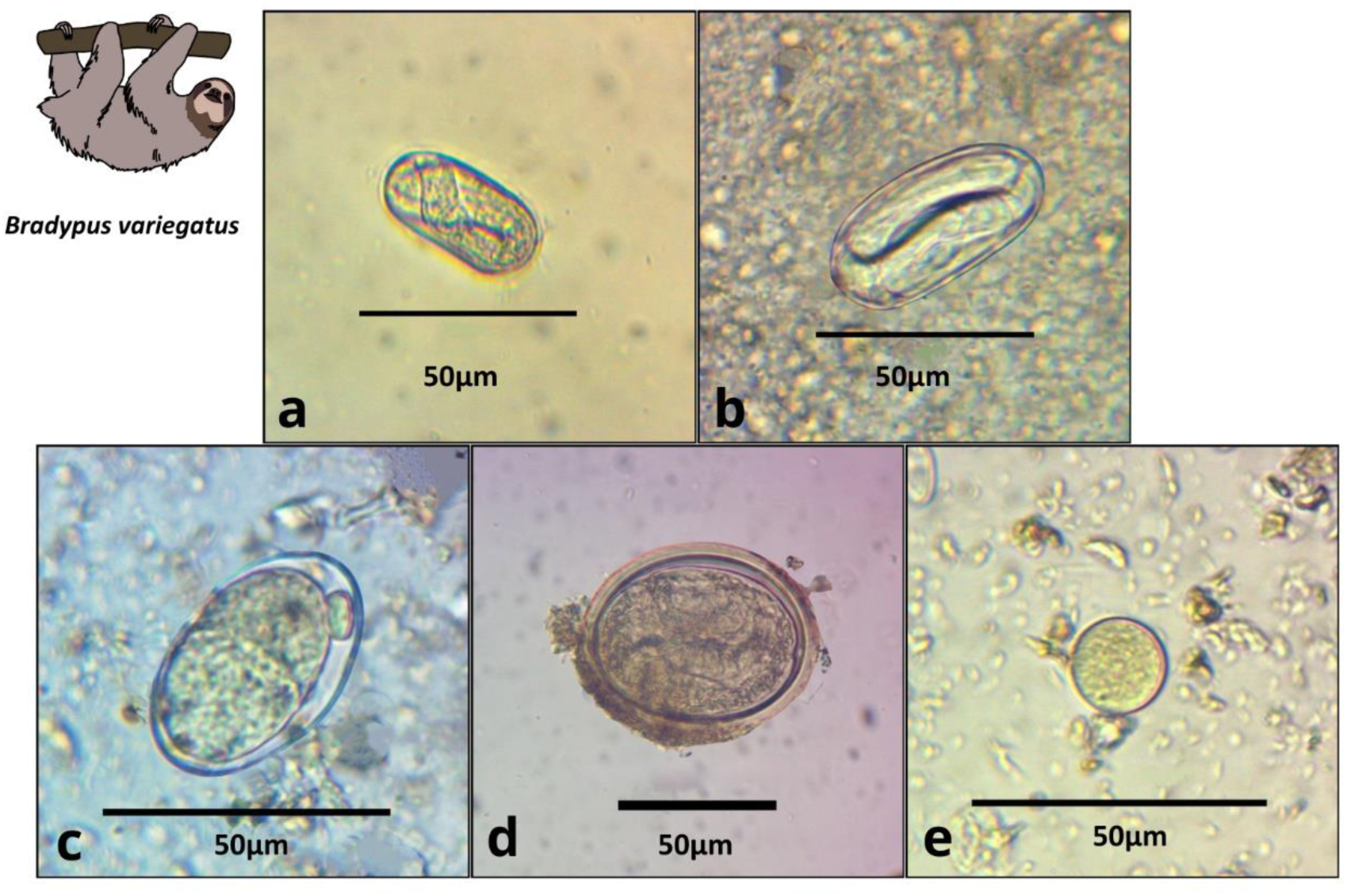
Images of parasite groups identified in *B. variegatus*. Figure shows morphology and size. **a)** Spirocercidae egg morphotype 1, **b)** Spirurida egg morphotype, **c)** Subuluridae egg morphotype, **e)** Ascaridida egg, e) Coccidia cyst morphotype 1.

**Figure 3.**
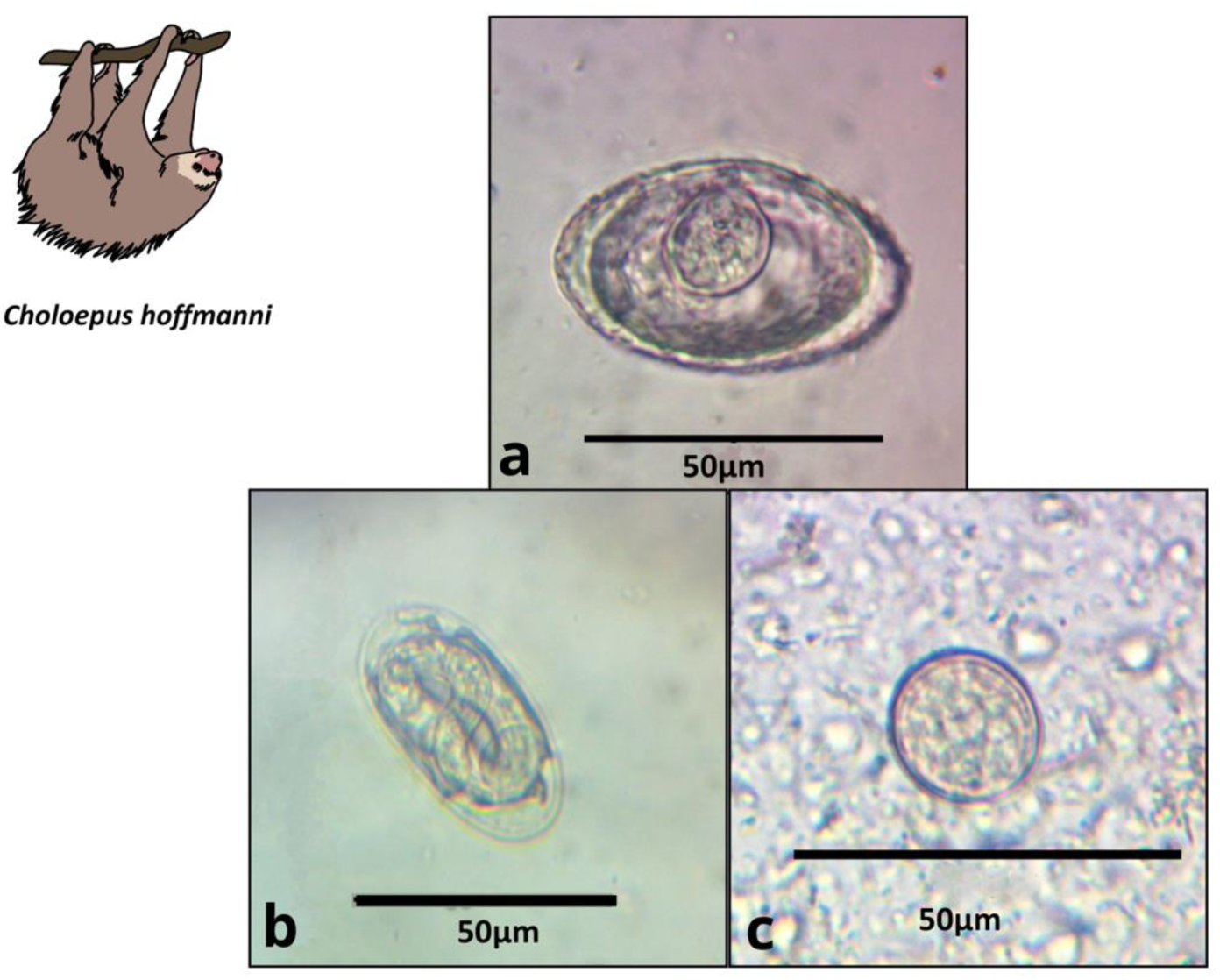
Images of parasite groups identified in *C. hoffmanni*. Figure shows morphology and size. **a)** Anoplocephalidae morphotype, **b)** Spirocercidae egg morphotype 2, **c)** Coccidia cyst morphotype 2.

**Table 1.** Parasitic taxa of sloth species. Eggs characterized as GI parasites were grouped into morphotypes and identified to the finest taxonomic level possible within 3 phyla and 5 families. Representative photos in Figures 2-3, mean length × width measurements (parentheses show number of eggs measured), shapes, general descriptions, and observed prevalences of each morphotype.

| Host | Egg morphotype | Phylum | Size ($\mu\text{m}$) | Shape | Description | Prevalence |
| --- | --- | --- | --- | --- | --- | --- |
| *Bradypus variegatus*<br>N = 22 | Spirocercidae egg morphotype 1<br>(Fig. 2a) | Nematoda | 38.8 × 18.6 (1) | Oval | Thin shell, larval eggs | 4.2% |
| | Spirurida egg morphotype (Fig. 2b) | Nematoda | 50.0 × 23.7 (12) | Oval | Thick, smooth shell, larval eggs | 27.3% |
| | Subuluroidea egg morphotype (Fig. 2c) | Nematoda | 40.5 × 24.1 (2) | Oval | Smooth and thin shell, no larval development | 4.2% |
| | Ascaridida egg morphotype (Fig. 2d) | Nematoda | 77.9 × 63.0 (2) | Rounded | Thick shell, larva visible inside, | 9.1 % |
| | Coccidia cyst morphotype 1 (Fig. 2e) | Apicomplexa | 25.4 × 23.4 (10) | Round | Thin shell | 31.9% |
| *Choloepus hoffmanni*<br>N = 16 | Anoplocephalidae morphotype 1<br>(Fig. 3a) | Platyhelminthes | 47.8 × 36.1 (12) | Triangular to oval | Thick wall, visible pyriform apparatus | 43.8% |
| | Spirocercidae egg morphotype 2<br>(Fig. 3b) | Nematoda | 49.0 × 23.6 (2) | Oval | Thin double shell | 6.3% |
| | Coccidia cyst morphotype 2 (Fig. 3c) | Apicomplexa | 22.3 × 18.7 (16) | Rounded | Thin shell | 56.3% |

### Statistical results

We found no significant difference in mean parasite richness between sloth species (ANOVA: *F*_1,34_ = 1.3, *P* = 0.263) or habitat types (*F*_1,34_ = 0.4, *P* = 0.544), but there was a marginally significant sloth species × habitat type interaction (*F*_1,34_ = 3.6, *P* = 0.066). This interaction reflected the elevated parasite richness in *C. hoffmanni* samples from undisturbed habitats (Figure 4). This effect of habitat was evidently strong in *C. hoffmanni*, albeit statistically underpowered, as the mean parasite richness was ∼2-fold greater in undisturbed habitats compared to disturbed habitats. In contrast to patterns of parasite richness across sloth species, the composition of parasites observed in *B. variegatus* differed profoundly from *C. hoffmanni*. There was a strong and statistically significant difference in parasite composition between sloth species (PERMANOVA: pseudo-*F*_1,21_ = 13.0, R^2^ = 0.37, *P* ≤ 0.001), but not between habitat (pseudo-*F*_1,21_ = 0.7, R^2^ = 0.02, *P* = 0.587), and there was no significant sloth species × habitat interaction (pseudo-*F*_1,21_ = 0.6, R^2^ = 0.02, *P* = 0.716). As we observed no shared parasites between sloth species, mean Jaccard dissimilarity between these host species was 1.00, far exceeding the mean differences among samples within species overall (*B. variegatus =* 0.71, *C. hoffmanni =* 0.46) and between habitats (disturbed vs. undisturbed populations of *B. variegatus =* 0.70, *C. hoffmanni =* 0.43; Table 1).

**Figure 4.**
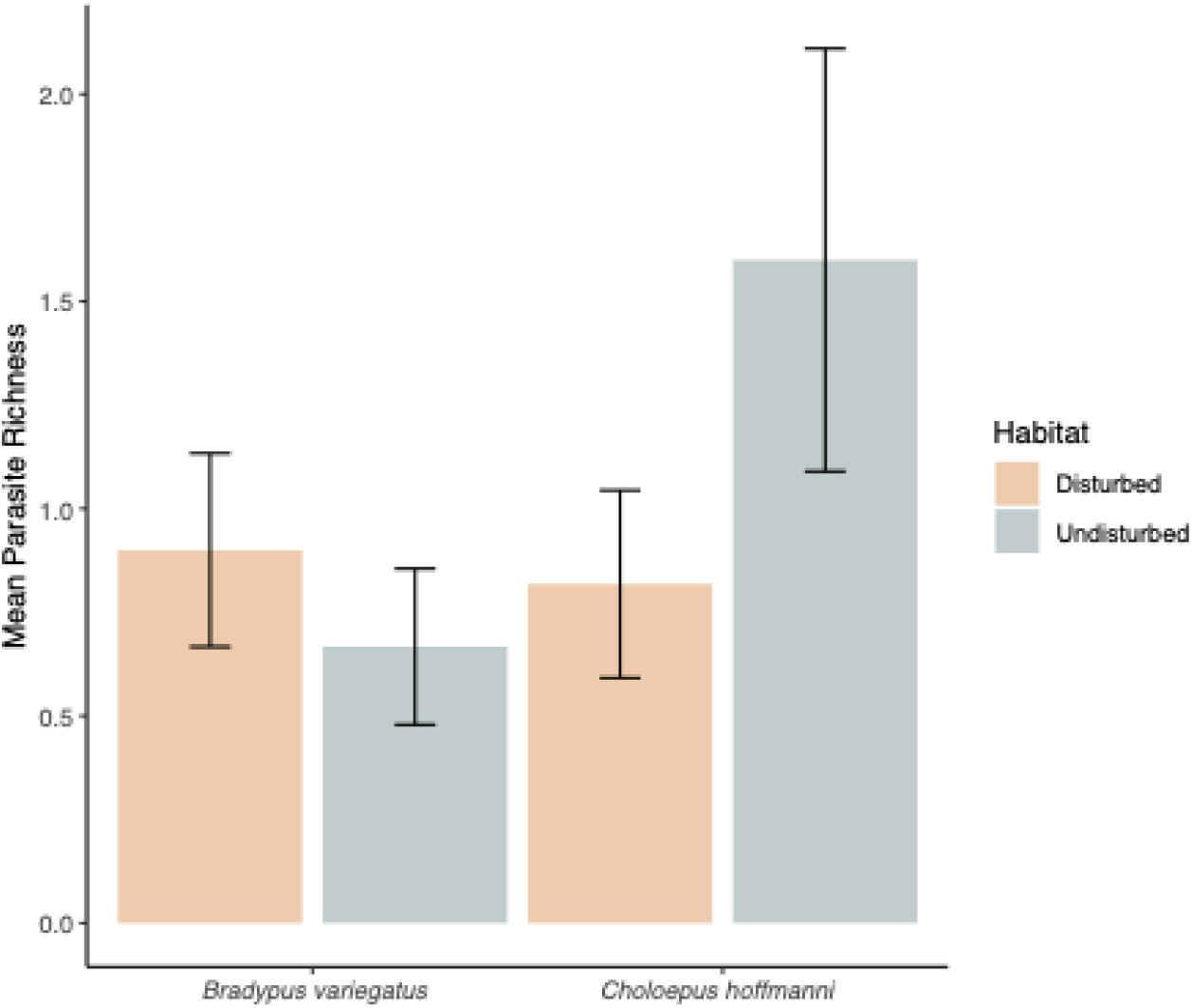
Mean parasite richness across sloth populations in each species-habitat combination. The bars represent standard error.

## Discussion

We asked whether sloth-parasite interactions are structured primarily by host identity or the local environment. Our comparisons of two co-occurring sloth species from regions characterized by different levels of human disturbance revealed five parasitic morphotypes from *B. variegatus* and three from *C. hoffmanni*. The parasite communities were mutually exclusive to sloth species, even between populations co-occurring in the same sites, suggesting a high degree of host specificity in these host-parasite interaction networks. Furthermore, many parasitic morphotypes were shared between conspecific populations that were geographically and ecologically separated across non-disturbed and disturbed habitats. Our results demonstrate a strong role for host species identity in structuring the composition of parasite assemblages in sloths.

Coproparasitology works well when the focus of a study is comparative because it allows for internal consistency of taxon identification. By contrast, it is notoriously difficult to compare across studies, and the best practice is to accept some imprecision in identifications when reviewing comparisons to the relevant literature (Akrim et al. 2018: Rojas et al. 2024). For each of the 8 morphotypes we identified across host species (Figures 2-3), we reviewed what was previously documented in sloths (Appendix S1). We identified 11 relevant papers involving both egg structures and adult parasites dating to 1928, including one published student theses. We found evidence that five of the taxa that we identified were similar to those previously reported in sloths, whereas we could find no evidence of the other three having been previously reported in association with sloths and conclude that they are being documented for the first time here (Appendix S1). Our goal was not to positively identify parasite species as our focus was on quantifying the degree of parasite overlap within and between species across these habitat types, but this review of the prior literature highlights the benefits of being conservative when drawing taxonomic inferences based on coproparasitological data and emphasizes the potential benefits for future studies (Appendix S1). This approach has been successfully implemented across a diverse array of wild animals, including primates, felines, carnivores, armadillos, and rodents (Kooriyama et al. 2012; Parr et al. 2013; Rondón et al. 2017; Agostini et al. 2018; Solórzano-García & de Leon 2017; Hou et al., 2020; Hewavithana et al. 2022; Bellusci et al. 2024).

Even with our conservative approach to the identification of parasitic morphotypes, our study revealed greater richness of sloth GI richness than all prior publications combined (Sibaja-Morales et al., 2009; Vicente et al., 1997). Our observations raise questions about how the sloths ingest such a diversity of parasites, especially since the predominant parasitic taxa that we found infecting both sloth species rely on intermediate hosts to complete their life cycles. We can identify at least three non-mutually exclusive hypotheses: (*i*) although documented almost exclusively in captivity, sloths are capable of omnivory, and they may intentionally scavenge for animal material on occasion, (*ii*) sloths incidentally ingest arthropods or their frass on vegetation while foraging (e.g., floral mites), or (*iii*) sloth-associated arthropods such as moths and beetles act as carriers from the stool to the host, and thus contaminating the fur and mouth of the sloth (Denegri et al., 1998; Madrigal, 2020; Wickström, 2004). We cannot differentiate between hypotheses *ii* or *iii*, but several lines of evidence suggest hypothesis *i* is insufficient to account for the prevalence of parasites that require intermediate hosts infecting both species. First, although it is known that *C. hoffmanni* can engage in omnivory because they accept eggs in captivity (Reyes-Amaya et al., 2015), and although prior studies of sloth diets have focused almost exclusively on herbivory (Sánchez-Chavez, 2021; Vaughan et al., 2007), and there is no evidence of intentional omnivory in *B. variegatu*s that could explain the diversity and prevalence of Spirocercidae infecting both species. Second, arthropods represent the most likely intermediate hosts of these parasitic taxa: common intermediate hosts of Spiruroidea include beetles and other arthropods (Bain et al., 2013; Chabaud & Bain, 1994), while oribatid mites or collembolans may often serve as intermediate hosts for Anoplocephalidae cestodes (Wickström, 2004). Sloth-associated beetles, mites, and moths live in sloth fur, for example, and some of these arthropods oviposit in sloth feces, creating ample opportunities to transfer infectious material to sloths for ingestion. Additional work will be needed to differentiate between hypotheses *ii* and *iii*, especially as both direct (e.g., arthopod carriers) and indirect (e.g., food-borne) routes of fecal-oral transmission are plausiblearthropods to serve as intermediate hosts of these GI parasites could help elucidate transmission pathways.

Our study is the first to report on the GI parasites of sloths in primary forest, reflecting the long-standing difficulty of obtaining samples from sloth populations in structurally complex habitats compared to more disturbed areas such as urban green spaces and agricultural areas. Although our sample set from the primary forest at La Selva was relatively small, it supports several important conclusions. First, there was no strong or significant effect of habitat on the composition of parasites infecting sloths. This is in part because we encountered so many undescribed helminth taxa in sloths from primary forest that we had a limited ability to discern whether these taxa comprise a community of parasites that is characteristic of primary forests or if they were simply rare and potentially opportunistic parasites of the sloths we encountered. Second, the mean parasite richness of *C. hoffmanni* was strongly, albeit marginally significantly, greater in primary forest than disturbed habitats—a trend driven largely by the elevated prevalence of cestodes in forest sloths (Figure 4). Third, although protozoa are commonly associated with elevated stress levels (Fayer, 1980), and although it would be reasonable to hypothesize that sloths experience elevated stress in human-dominated environments, we did not observe elevated protozoa of the family Eimeriidae (Apicomplexa) infection rates in urban sloths (41% of 17 primary forest samples vs. 43% of 21 urban samples). These outcomes could reflect differences in the abundance and diversity of intermediate hosts required in parasite lifecycles, differences in the timing or extent to which GI infections spread within and across geographically discrete populations, and/or differences in the transmissibility of each parasite among hosts according to habitat—especially since sloth diets, movements and foraging behaviors, and population densities are expected to vary as a function of habitat type (Pool et al., 2016; Silva et al., 2013; Werner & Nunn, 2020).

Knowledge gaps concerning sloth parasites could be addressed in several ways. First, given the high diversity of novel parasites we observed, and given the difficulty of precisely identifying parasites when observing their eggs, necropsies to collect and properly identify adult parasites would enable more comprehensive efforts to compare the parasites of free-ranging sloths. Studies of *Bradypus* spp. in Brazil, for example, have characterized five adult gastrointestinal nematode species (Michel et al., 2017; Santos & Werneck, 2013; Werneck et al., 2008). A lack of adult nematode specimens that can be positively identified leaves us uncertain if the parasites infecting *Bradypus* in Costa Rica are the same as have been documented in Brazil. Second, evaluations of how GI parasites affect sloth health would require veterinary assessment and may be a function of both parasite diversity and load. Combinations of methods including fecal DNA analysis and egg counts could help better characterize the abundance, diversity, and potential health risks posed by parasites throughout the range of sloths (Chan et al., 2022; Titcomb et al., 2022). Third, whereas our study highlighted the diversity of sloth parasites that may be discovered in primary forests, our comparisons were based on just two geographically discrete habitats and further comparative studies could help clarify or reinforce the differences in parasite diversity that we observed here.

Lastly, sloths are among the species most frequently admitted to veterinary clinics for treatment and release following encounters with domestic wildlife, electrocution, or poor health. Although both species are listed as Least Concern (LC) on the IUCN red list (Moraes-Barros et al., 2022; Plese et al., 2022), some suggest that *C. hoffmanni* would be reclassified as threatened if more data were available (Sánchez-Chavez, 2021). Considering these many newfound sloth-parasite interactions and the unknown health effects they may have, caution to minimize the risk of transmitting harmful parasites between populations and species may be warranted when undertaking ecological rewilding programs, translocations, or *ex situ* care of wild sloths.

## Declarations

## Acknowledgements

We thank Toucan Ranch, Kids Saving the Rainforest, the Sloth Sanctuary, and the Organization for Tropical Studies for facilitating our research. We thank the government of Costa Rica and SINAC for permission to conduct this research.

## Ethical approval

No animal in this study was subjected to experimentation or manipulation.

## Consent to participate

All the authors consented to participate in this study.

## Consent for publication

All the authors consent to publication of this article.

## Availability of data and material

All data are reported in tables in the main text and supporting materials.

## Author contributions

Conceptualization and project design: EAV, JN, TRK. Fieldwork: EAV, MGF, JG, RC, TRK. Laboratory analysis EAV, MGF. Statistical analysis EAV, MGF, TRK. Manuscript writing and revision EAV, MGF, JN, RC, TRK.

## Appendix S1

Studies of gastrointestinal parasites from sloths are limited, but nevertheless prior publications provide a basis for interpretation of our results. We reviewed all publications known to describe taxonomic identifications of gastrointestinal parasites from sloths, and found that reports were limited to only two host species of the genus *Bradypus*, specifically *B. variegatus* and *B. tridactylus*. There have been no prior descriptions of adult parasites from any of five the remaining sloth species (*Bradypus torquatus*, *B. crinitus*, *B. pygmaeus*, *Choloepus hoffmanni*, or *Choloepus didactylus*). In publications dating to 1928, a total of eight nematode species, one cestode, and one protozoan have been described in association with three-fingered sloths (Table S1). Coproparasitological studies are also limited, although two prior studies of sloths occurring in captivity and rural areas have reported observations of nematode, cestode, and protozoan eggs (Lainson & Shaw 1982; Sibaja-Morales et al., 2009). As with the current study, taxonomic identifications of potential parasite species based on observations of eggs can be challenging and thus the ability of these descriptions to support cross-study comparisons is limited.

Here, we provide a detailed review of taxonomic and morphological similarities between our results and the available literature. We err on the side of ensuring broadly accurate taxonomic descriptions, even in cases where taxonomic resolution is limited.

Five parasite morphotypes from *Bradypus variegatus* (Table 1; Figure 2).

1. Spirocercidae egg morphotype 1 (Figure 2a): This morphotype was found in one sample, and it resembles the Spirurida egg morphotype (Figure 2b), but its size is significantly smaller (Table 1). The family Spirocercidae has been reported in *Bradypus variegatus* and *B. tridactylus*, with *Leiurus* and *Paraleiurus* being the only described genera (Table S1). Egg sizes vary between the suite of host and Spirocercidae species found in sloths: *Paraleiurus vazi* (Vicente & Gomez, 1971) and *Leiurus pereirai* (Gomez & Vicente, 1970) exhibit similar measurements and larvated eggs.
2. Spirurida egg morphotype (Figure 2b): This morphotype was the most prevalent nematode egg in *B. variegatus* (Table 1). These eggs could potentially match prior descriptions of *Leiurus leptocephalus*, as adult worms from this species were reported in both Costa Rica and Brazil (Jiménez-Quirós & Brenes, 1956; Werneck et al., 2005) (Table S1). Description of these eggs are based on gravid female worms included measurements of 56 by 22 μm (Vaz & Pereira, 1929), 33 by 13 μm (Jiménez-Quirós & Brenes, 1956), and an average of 44 by 24 μm (Werneck et al., 2005). This variability in egg measurements may be attributable to differences in female reproductive maturity, host species, and/or geographic variation.
3. Subuluridae egg morphotype (Figure 2c): This morphotype is being reported for the first time in *B. variegatus*. No prior publication of this family in association with sloths were identified in our review.
4. Ascaridida egg (Figure 2d): This morphotype is being reported for the first time in *B. variegatus*. No prior publication of this family in association with sloths were identified in our review.
5. Coccidia cyst morphotype (Figure 2e): Cysts of the protozoan Eimeridae (Apicomplexa) were reported in prior coprological surveys in sloth species (Lainson & Shaw 1982; Sibaja-Morales et al., 2009). It is not practical to provide a precise identification of protozoan species based on observations of cysts, further studies using specialized Protozoa techniques would be required to provide better taxonomic resolution.

Three parasite morphotypes from *Choloepus hoffmanni* (Table 1; Figure 3).

1. Anoplocephalidae morphotype (Figure 3a): This was the most prevalent helminth egg morphotype detected (Table 1). This family of Cestoda was reported in other coproparasitological studies in Costa Rica, though none reported egg sizes (Sibaja-Morales et al., 2009; Estrada-Rodríguez, 2007). The only Cestoda that has been described in sloths is *Moniezia benedini*, which was detected in *B. variegatus* (Flores-Barroeta et al., 1958).
2. Spirocercidae egg morphotype 2 (Figure 3b): This morphotype is being reported for the first time in *Choloepus* spp. No prior publication of this family in association with sloths were identified in our review.
3. Coccidia cyst morphotype 2 (Figure 3c): Cysts of the protozoan subfamily Eimeridae were reported in prior coprological surveys in sloths species (Lainson & Shaw 1982;; Sibaja-Morales et al., 2009). It is not practical to provide a precise identification of protozoan species based on observations of cysts, further studies using specialized Protozoa techniques would be required to provide better taxonomic resolution.

Coproparasitological studies on free-ranging sloths that provide morphological characterizations of parasitic eggs and include images have served as key points of reference for our observations. In our review, we identified several inaccuracies in prior studies that are frequently cited in the relevant literature (Estrada, 2007; Sibaja-Morales et al., 2009). Estrada’s (2007) undergraduate thesis on sloths in cacao plantations and Sibaja-Morales’ (2009) peer-reviewed paper on captive sloths both report *Leiurus* and *Moniezia* specimens, but they lack illustrations, measurements, or voucher specimens for verification. The limited available data reflects the difficulty in assigning species to microscopic eggs and underscores the need for more robust methodologies that is not exclusive to sloths. Therefore, rather than assigning precise species or identifications that extend beyond the resolution of our data, we emphasize the need for further research using complementary techniques such as necropsies, systematic parasite collections, and molecular analyses to better identify wildlife parasites.

**Table S1.** Gastrointestinal parasite species that have been identified based on adult specimens in association with sloths in prior publications.

| Host | Parasite species | Free or captive | Country | Citation |
| --- | --- | --- | --- | --- |
| ***Bradypus tridactylus*** | *Leiuris leptocephalus* (Nematoda) | Free ranging | Brazil | Vaz & Pereira 1929 |
|  | *Leiuris pereirai* (Nematoda) | Free ranging | Brazil | Gomes & Vicente 1970 |
|  | *Leiuris vazipereirai* (Nematoda) | Free ranging | Brazil | Vaz & Pereira 1929 |
|  | *Paraleiuris locchii* (Nematoda) | Free ranging | Brazil | Vaz & Pereira 1929 |
|  | *Paraleiuris vazi* (Nematoda) | Free ranging | Brazil | Vicente & Gomes 1971 |
|  | *Giardia duodenalis* (Protozoa) | Captive | Brazil | Dos Reis et al., 2023 |
| ***Bradypus variegatus*** | *Leiuris leptocephalus* (Nematoda) | Free ranging | Costa Rica | Jiménez-Quirós & Brenes 1956 |
|  |  | Free ranging | Brazil | Werneck et al., 2008 |
|  | *Paraleiuris vazi* (Nematoda) | Captive | Brazil | Araujo et al., 2021 |
|  | *Paraleiuris locchii* (Nematoda) | Captive | Brazil | Michel et al., 2017 |
|  | *Moniezia benedeni* (Cestoda) | Free ranging | Costa Rica | Flores Barroeta et al., 1958 |
|  | *Giardia duodenalis* (Nematoda) | Captive | Panama | Hegner & Schumaker 1928 |

